# Cold adapted desiccation-tolerant bacteria isolated from polar soils presenting high resistance to anhydrobiosis

**DOI:** 10.1101/2021.02.06.430066

**Authors:** Felipe Nóbrega, Rubens T. D. Duarte, Adriana M. Torres-Ballesteros, Luciano Lopes Queiroz, Lyle G. Whyte, Vivian H. Pellizari

**Author notes:** **Correspondence**: Felipe Nóbrega, Oceanographic institute, University of São Paulo, Praca do Oceanografico, 191, São Paulo – SP – Brasil – 05509-900.

## Abstract

Life on Earth is strictly dependent on liquid water. In polar terrestrial environments, water exists in solid state during almost the entire year. Polar microorganisms have not only to adjust their metabolism to survive at subzero temperatures, but also need to cope with extremely dry conditions. We investigated the presence of desiccation-adapted bacteria in Arctic permafrost and Antarctic surface soils and characterized their survivability to dryness. We selected desiccation tolerant cells by treating the soils with chloroform prior to cultivation, in order to mimic the stress of low water activity for long periods. From over 1000 colonies from different samples, 23 unique strains were selected and identified as members of phyla *Firmicutes, Proteobacteria* and *Actinobacteria*. About 60% of the strains survived after 50 days in anhydrobiosis. The competence to withstand desiccation varied between close related strains isolated from different locations, bringing the question if environmental conditions may play a role in the observed desiccation tolerance. Survivability was also affected by the solution in which the cells were suspended before drying; R2B medium being more protective than water. This is the first time that chloroform was used to select desiccation tolerant microorganisms from polar soils. The collection of polar microorganisms described herein opens the possibility of further experiments aiming to investigate the resistance mechanisms of polar anhydrobionts. Desiccation tolerance is fundamental to the survivability of microorganisms to the space environment and at the surface of thin-atmosphere planets like Mars. Therefore, the selected strains may open a road to better understand the limits of cold adapted life on Earth and beyond, and compare mechanisms of resistance with anhydrobionts from divergent extreme environments.

## Introduction

Life as we know it cannot thrive without water; it can survive through dry seasons (Stevenson et al., 2015; Chaplin, 2006). In polar terrestrial environments, water exists in solid state during almost the entire year. Still, microbial communities in these regions are diverse, even in less dynamic ecosystems such as the permafrost (Steven et al., 2006; Jansson and Tas, 2014). This state of water stress is known as anhydrobiosis – a state of extreme metabolism reduction and physiological adaptations (Billi and Potts, 2002). The study of the physiology of anhydrobionts (one of the terms used to address desiccation tolerant species) depends on strains isolation. It came to our attention a culture method for isolation of desiccation resistant microorganisms, from soils subjected to seasonal droughts, using chloroform (Narvaez-Reinaldo et al., 2010). This methodology exploits the effect of bacterial protective measures taken during anhydrobiosis that not only prevent damage related to desiccation, but also to exposure to organic solvents (Manzanera et al., 2004; Vilchez et al., 2008). It is unknown if microorganisms endemic to cold regions have the same drought tolerance and survival mechanisms like other well-studied anhydrobionts; and whether this would be a valid method for isolation of polar microorganisms. Experiments using desiccated microorganisms vary, not only regarding the experimented strains, but also in the solution that the cells are suspended before drying. Commonly used solutions are saline solutions of different concentrations, culture medium and water. It is not usually addressed in these experiments the effect of the drying solution in bacteria survivability, although protective solutions can induce desiccation resistance (Julca et al., 2012).

The understanding of the limits for cellular life on Earth and beyond – including water absence – is under the realm of astrobiology. Experiments performed at the International Space Station and in simulation chambers on Earth, investigate the effects of environmental factors found in outer space (e.g. pressure, desiccation, temperature, UV and radiation) in algae, lichens and bacteria (Horneck et al., 2001; Mancinelli, 2015; Rabbow et al., 2009; Olsson-Francis and Cockell, 2010; Meessen et al., 2015). In most, if not all, of these experiments, desiccation and low temperature tolerance preconditions microbial survival – which underscores the importance of studying cold adapted anhydrobionts. The likelihood of extant microbes in comets or meteorites (central to the Panspermia hypothesis) is also dependent on life tolerance to extremely dry and cold conditions (Clark, 2001). On Mars, microbial life is still a possibility; its subsurface soil contains water ice, and liquid water (if only ephemeral) any microbial life that might exist must cope with desiccation (Boynton et al., 2002; Ojha et al., 2015).

This study aims were: (1) to test an isolation methodology for desiccation resistant polar microorganisms (from soil samples from the Arctic and Antarctica); (2) to assess each strain’s desiccation tolerance over time (up to 50 days) and the drying solution influence (water vs. culture medium R2B 10%); (3) to investigate the relationship between strain’s phylogeny and its habitat to the measured desiccation tolerance.

## Materials and Methods

### Soil samples

Soil samples were collected from two regions in Antarctica – soil exposed by glacier retreat (Baranowski Glacier, King George Island) and volcanic soil from Deception Island – and from one region in the Canadian High Arctic (Table 1). Sampling in Antarctica was made during 2008/2009 summer; sampling in the Arctic was made during July 2010. Antarctic soils were collected from the surface (no more than 10 cm deep) with sterile materials and kept frozen (−20 °C to -50 °C) in sterile plastic bags until usage in 2013. Arctic sample were collected on polygonal permafrost terrain located near the McGill Arctic Research Station (MARS); a core was taken as previously described and kept frozen until usage (Wilhelm et al., 2012), and from it three soil depths were analyzed (Table 1).

**Table 1.** Soil samples used for the isolation of desiccation resistant microorganisms.

| Sample | Description | Sampling sites | Coordinates |
| --- | --- | --- | --- |
| Dec85 | Border of Deception's caldera (5 °C) | Deception Island | 62°55'21.55"S |
|  |  | Antarctica | 60°38'7.19"O |
| Dec92 | Border of geothermal lake (10 °C) | Deception Island | 62°58'50.00"S |
|  |  | Antarctica | 60°34'24.00"O |
| Dec94 | Border of geothermal lake (6 °C) | Deception Island | 62°58'46.31"S |
|  |  | Antarctica | 60°34'31.65"O |
| Bas01 | Distance from glacier: 0 m | Baranowsky | 62°11'53.42"S |
|  |  | Antarctica | 58°26'53.19"O |
| Bas06 | Distance from glacier: 189 m | Baranowsky | 62°11'52.87"S |
|  |  | Antarctica | 58°26'40.19"O |
| Bas12 | Distance from glacier: 473 m | Baranowsky | 62°11'52.56"S |
|  |  | Antarctica | 58°26'20.62"O |
| ArcA | Active Layer - Depth: 25 cm | Axel Heiberg | 79°24'57.90"N |
|  |  | Arctic | 90°45'48.12"O |
| ArcM | Transient Layer - Depth: 50 cm | Axel Heiberg | 79°24'57.90"N |
|  |  | Arctic | 90°45'48.12"O |
| ArcP | Permafrost - Depth: 100 cm | Axel Heiberg | 79°24'57.90"N |
|  |  | Arctic | 90°45'48.12"O |

### Strain isolation

In order to mimic the selective pressure exerted by the absence of water for a prolonged time, 5 grams of air-dried soil was incubated with 10 mL of ≥ 99.8% chloroform for 30 minutes at room temperature in sterile conditions, the first step of our isolation and selection of desiccation-tolerant bacteria (Fig. 1). This step was performed according to the methodology established by Narvaez-Reinaldo et al. (2005) with minor changes. After incubation, samples were air dried for 30 minutes until complete chloroform evaporation. Treated soils were then stirred (not shaken) with 10 mL R2B medium (EMD Chemicals Inc.) for 30 minutes. A total of 100 μL of the resulting solution was inoculated in R2A and TSA media (Difco Laboratories), and incubated at 30 °C and 0 °C until colony appearance.

**Figure 1.**
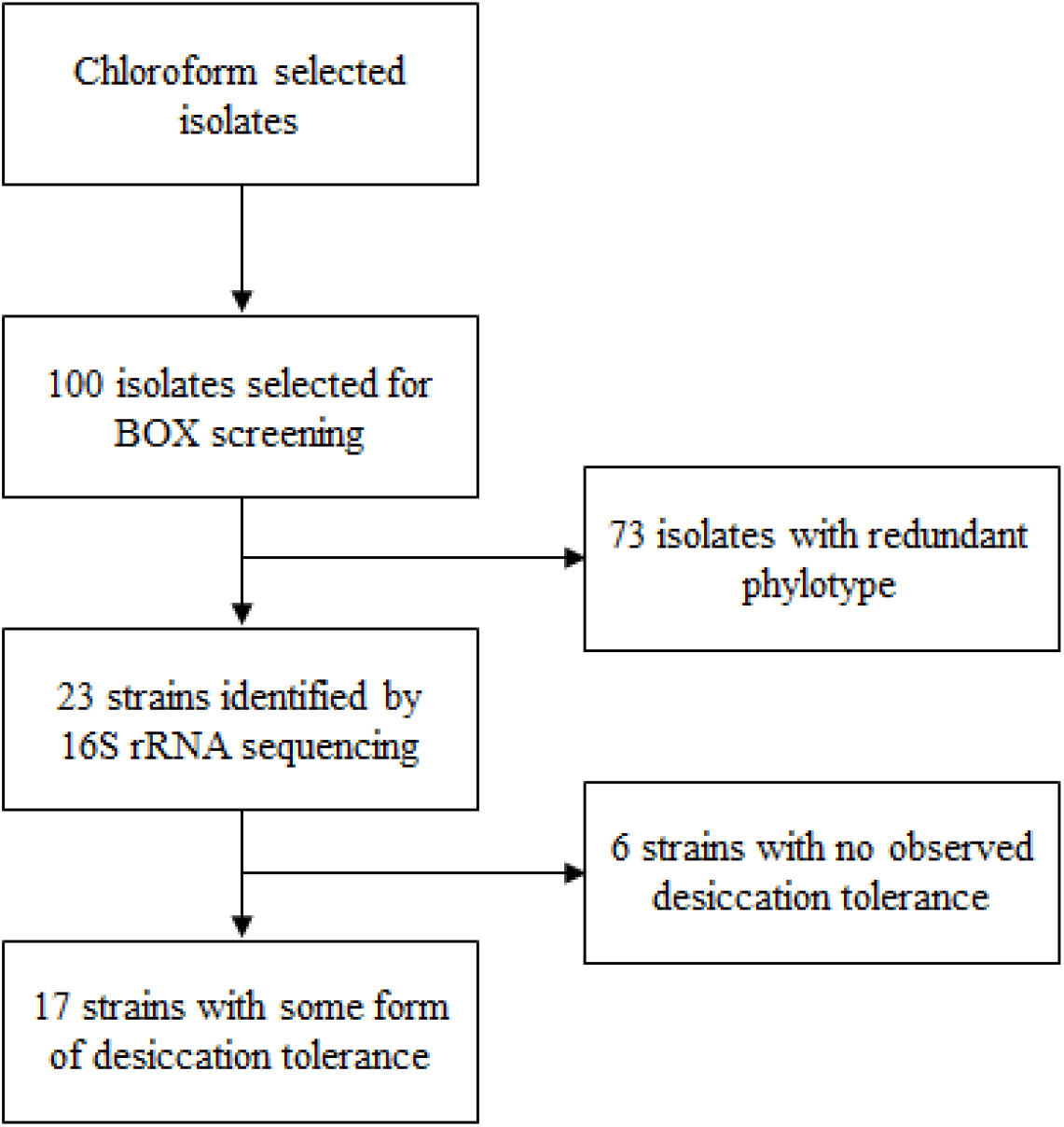
Strain selection diagram.

### Isolates identification

Aiming to select strains with unique phylotypes, we used the BOX-PCR fingerprinting technique, excluding the isolates with similar DNA amplification profile, as previously described (Martin et al., 1992). With the formerly selected isolates, part of the 16S rRNA gene was amplified for phylogenetic study, using the forward primer 27F (5’ AGAGTTTGATCMTGGCTCAG 3’) and the reverse primer 1492R (5’ TACGGYTACCTTGTTACGACTT 3’), following standard PCR methods (Goodfellow and Stackebrandt, 1991). Sequencing reaction was made with the primer 27F, yielding sequences with 700pb length. Sequences were submitted to the GenBank genetic sequence database (Submission ID: SUB1160085). Sequence quality was verified by Phred software (Ewing et al., 1998) and aligned against the SILVA database, using the SINA Web-aligner. The alignment was combined with 30 neighboring sequences from SILVA database using the ARB MERGE tool (Pruesse et al., 2012). ARB was used for reconstruction of Maximum Likelihood (ML) phylogenetic trees (Ludwig et al., 2004) with RAxML (Randomized Accelerated Maximum Likelihood), a random starting tree and GTR+C model. Support values for internal branches of the ML tree were obtained by bootstrapping (1000 replicates).

Isolates nomenclature carries information of its soil of origin (e.g. **<u>ArcA</u>**30R01), its isolation temperature (e.g. ArcA**<u>30</u>**R01), culture medium (ArcA30**<u>R</u>**01) and a differentiation number (e.g. ArcA30R01, ArcA30R02, etc.).

### Desiccation experiments

Isolates were grown in TSA or R2A plates, at room temperature, until colony formation up to 4 weeks. Two variables were tested; desiccation solution and desiccation time.

#### Desiccation solution

One colony of approximately 1 mm^2^ was suspended in 100 μL of ultrapure water or R2B medium 10% (Proteose Peptone: 0.5 g/L; Glucose: 0.5 g/L; Magnesium Sulfate Heptahydrate: 0.1 g/L; Sodium Pyruvate: 0.3 g/L; Yeast Extract: 0.5 g/L; Casein Acid Hydrolysate: 0.5 g/L; Soluble Starch: 0.5 g/L; Dipotassium Phosphate: 0.3 g/L). Two μL of the cell solution was then deposited on a sterile circular glass slide (5 mm^2^) and air-dried at room temperature (Relative humidity = 45 ± 4%). The untreated control was not allowed to dry.

#### Desiccation Time

Desiccation tolerance was examined by incubating the previously desiccated cells in a silica-gel exsiccator (Relative humidity = 5% ± 1%) for 0, 1, 5 and 50 days.

### Survival rate determination

After the desiccation treatment, dry cells were suspended thoroughly with 10 μL of their correspondent desiccation solution (ultrapure water or R2B 10%), plated in TSA or R2A and incubated at room temperature in sterile conditions until colony appearance. Ten-fold serial dilutions were made to obtain a manageable number of colony forming units per plate (30 ≤ CFU ≤ 300 per plate). Plates were made in triplicates for each dilution; we calculated the data mean and the standard deviation (SD). Strain survivability was determined from the ratio N / N0, where: N = Colony-forming units (CFU) after desiccation treatment; N0 = Control group, CFU without treatment.

*Deinococcus radiodurans* R1 (Anderson et al., 1956) and *Escherichia coli* K-12 strain MG1655 (Blattner et al., 1997) were used as a positive and negative control for desiccation tolerance, respectively. In addition, strains from Antarctica – *Exiguobacterium antarcticum B7* (Carneiro et al., 2012) – and the Arctic – *Planococcus halocryophilus* OR1 (Mykytczuk et al., 2012) *–* were also tested. All strains described in this study will be made available upon request. Survival rate comparisons of each isolate were done using Kruskal-Wallis test (p<0.05, Bonferroni correction).

## Results

### Isolation and phylogenetic study of strains

Colonies grew from all chloroform treated soil samples, except from two soil samples from Antarctica. We isolated strains from all incubation temperatures and culture media from the Arctic soils, but not from Antarctica. For example, from Deception Island, a region of volcanic activity, we only obtained colonies at 30 °C and TSA medium. From Antarctic soils exposed by the Baranowski glacier retreat, colonies grew at 30 °C and 5 °C, but only in TSA. We isolated around 1000 strains from all sites; CFU/g of the treated soil samples ranged between 10^2^ and 10^3^ (Table 2).

**Table 2.** Number of Colony Forming Units per gram of soil (CFU.g-1) isolated from chloroform treated soil samples.

| Sample | TSA 5 °C | TSA 30 °C | R2A 5 °C | R2A 30 °C |
| --- | --- | --- | --- | --- |
| Dec85 | 0 | 33 | 0 | 0 |
| Dec92 | 0 | 0 | 0 | 0 |
| Dec94 | 0 | 606 | 0 | 0 |
| Bas01 | 0 | 472 | 0 | 0 |
| Bas06 | 0 | 0 | 0 | 0 |
| Bas12 | 568 | 0 | 0 | 0 |
| ArcA | 173 | 126 | 13 | 40 |
| ArcM | 693 | 186 | 66 | 126 |
| ArcP | 543 | 320 | 53 | 53 |

We selected around 100 isolates for BOX amplification screening, 23 of which were selected by their unique BOX-PCR amplification profile for identification by partial 16S rRNA gene sequencing. Strains were part of the Phylum: Actinobacteria (1strain, Deception Island), Proteobacteria (4 strains, Deception Island) and Firmicutes (18 strains from all the sampling sites) (Fig. 2).

**Figure 2.**
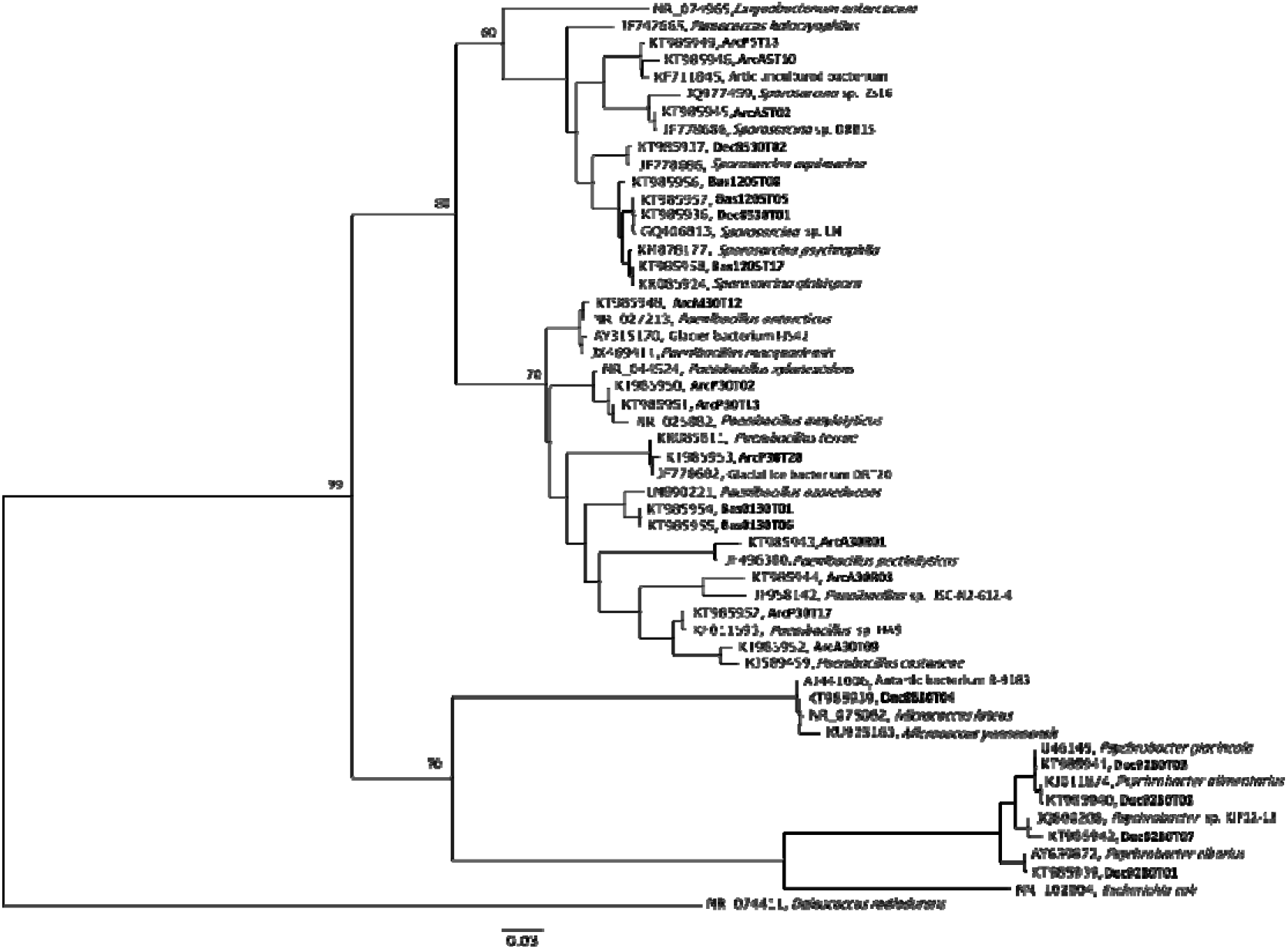
Phylogenetic tree based on gene 16S rRNA sequences showing the relationship between the desiccation-resistant isolates. The scale bar indicates 0.03 substitutions per nucleotide position. Bootstrap values (%), higher than 60 are shown in the nodes.

### Influence of suspension solution in desiccation tolerance

Six strains did not show resistance to desiccation under any circumstance: ArcP0513, ArcA05T10, ArcA05T02 – strains closely related to the *Sporosarcina* genus; ArcA30R03, ArcA30T09, ArcM30T12 – of the *Paenibacillus* genus. All strains presented a significantly higher survival rate when desiccated at R2B medium 10% (Fig. 3 and 4). Strains isolated from Deception Island were highly sensitive when desiccated in water suspension. Similarly, reference strains only showed desiccation tolerance when desiccated at R2B medium 10% with the exception of *D. radiodurans* (Fig. 3 and 4).

**Figure 3.**
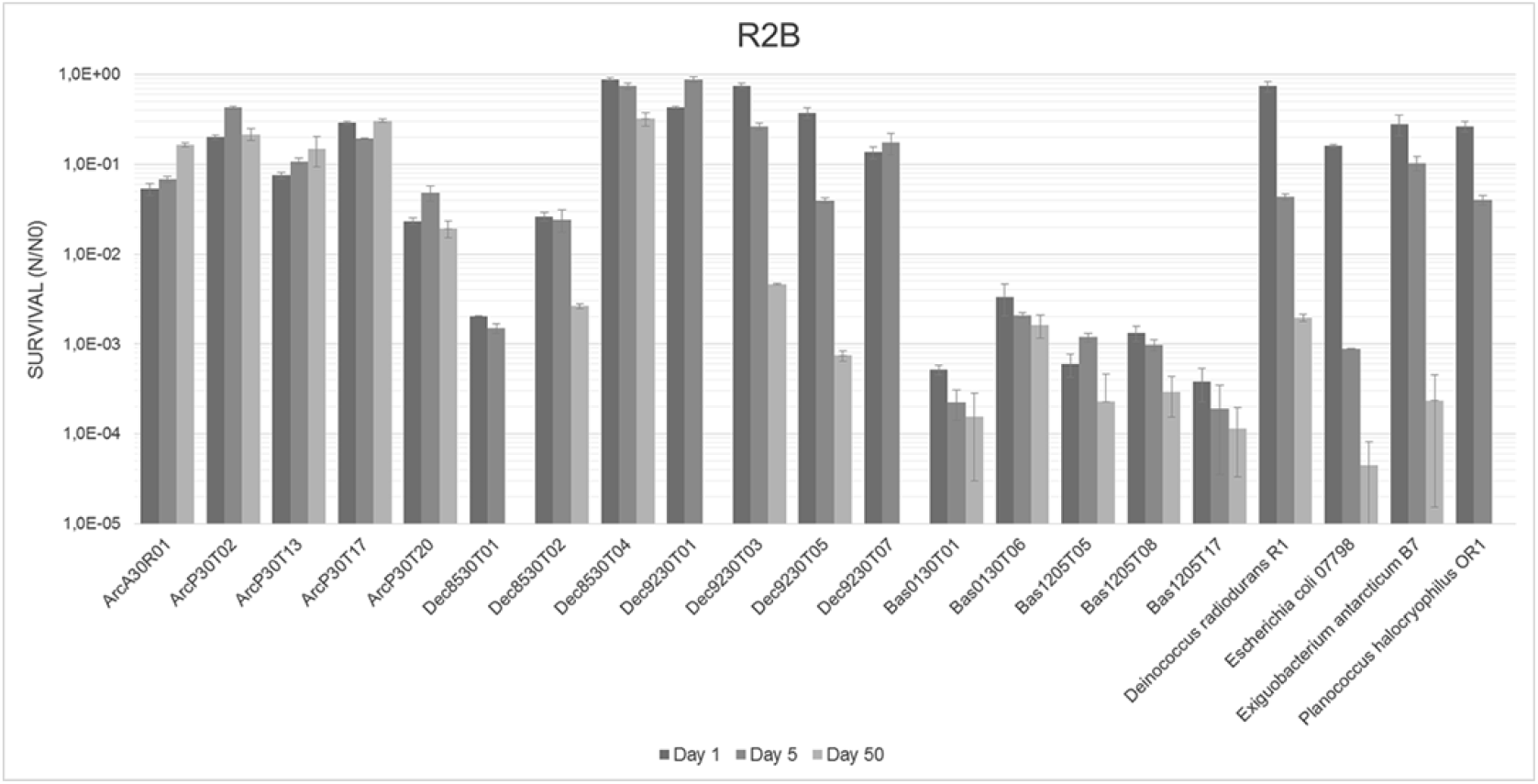
Survival of isolates after 1, 5 and 50 days of desiccation. One colony was suspended in a solution of R2B 10% medium, deposited on a glass slide and desiccated in an exsiccator. After the treatment, we suspended the cells in R2B 10% medium. Error bars show the standard deviations for three replicates.

**Figure 4.**
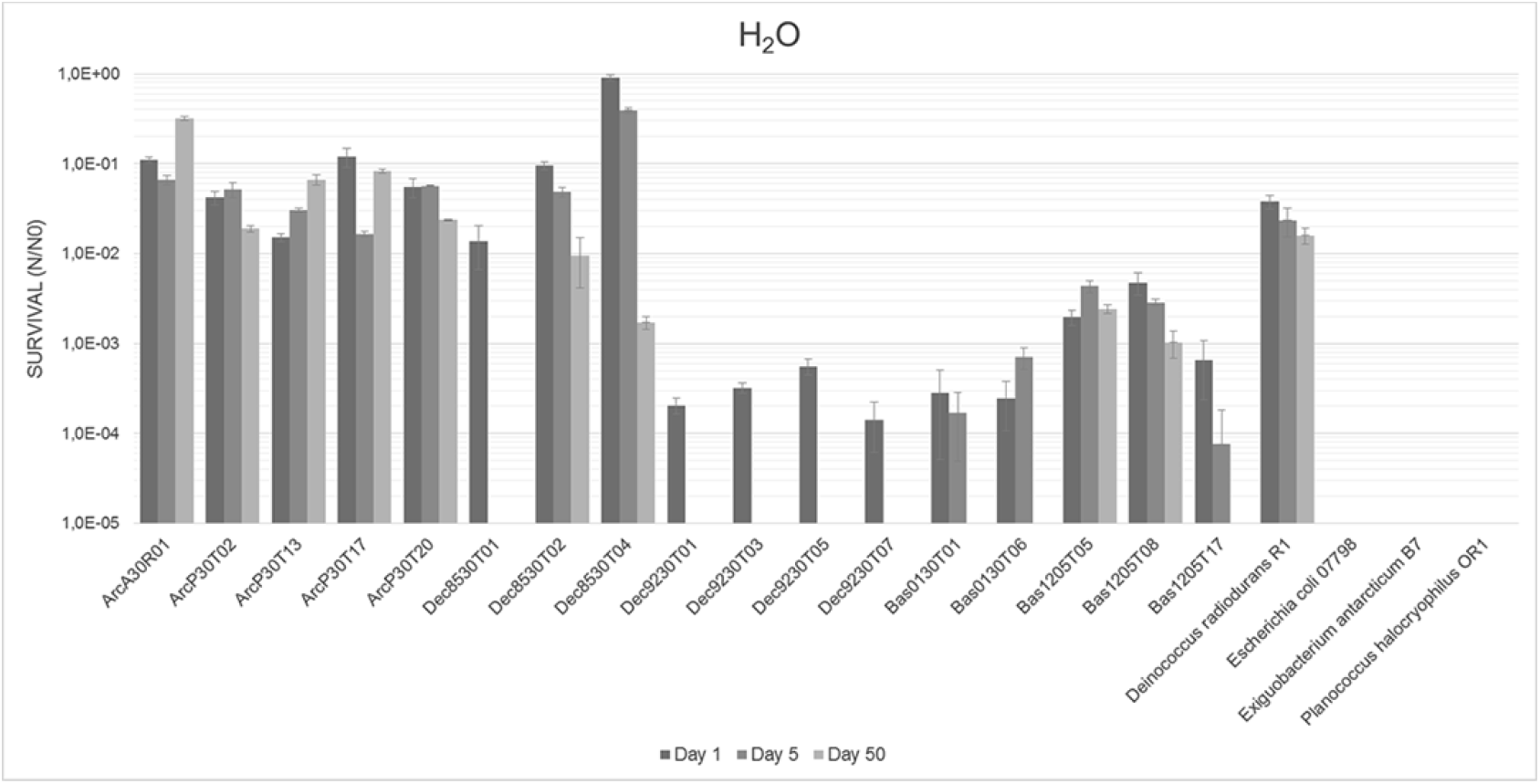
Survival of isolates after 1, 5 and 50 days of desiccation. One colony was suspended in a solution of ultrapure water, deposited on a glass slide and desiccated in an exsiccator. After treatment, we suspended the cells in H_2_O. Error bars show the standard deviations for three replicates.

### Isolates survival after different desiccation times

Most bacteria showed a reduction in survival rates with time (Fig. 3 and 4). Three strains desiccated in R2B 10% – ArcP30T02, ArcP30T17 and Dec8530T04 – were more tolerant to desiccation than *D. radiodurans*, showing less than a 10-fold reduction in viability after 50 days (Fig 3). *E. antarcticum* and *P. halocryophilus*, which suffered a 10-to 100-fold viability reduction after 5 days; after 50 days survival rates were similar to *E. coli*.

### Desiccation tolerance within bacterial families

Desiccation tolerance was not specific to a particular group (Table 3). Nevertheless, we found that the most resistant strains were within the Family *Micrococcaceae, Planococcaceae* and *Paenibacillaceae*. Gram-negative bacteria (*Moraxellaceae*) were more susceptible to desiccation, especially when desiccated in water suspension. Interestingly, strains from the same genera *Paenibacillus* and *Sporosarcina*, but isolated from different samples, showed different desiccation tolerance (Table 3).

**Table 3.** Summary of strain tolerance to desiccation for each phylum and family.

| Phylum | Family | Strain | Survival rate H<sub>2</sub>O |  |  |  | Survival rate R2B |  |  |  |
| --- | --- | --- | --- | --- | --- | --- | --- | --- | --- | --- |
|  |  |  | 1 day | 5 days | 50 days | Signif. | 1 day | 5 days | 50 days | Signif. |
| Firmicutes | Planococcaceae | ArcP5T13 | 0 | 0 | 0 | - | 0 | 0 | 0 | - |
|  |  | ArcA5T10 | 0 | 0 | 0 | - | 0 | 0 | 0 | - |
|  |  | ArcA5T02 | 0 | 0 | 0 | - | 0 | 0 | 0 | - |
|  |  | Dec8530T02 | 9,6E-02 | 4,8E-02 | 9,5E-03 | * | 2,6E-02 | 2,4E-02 | 2,6E-03 | ns |
|  |  | Bas1205T08 | 4,8E-03 | 2,9E-03 | 1,0E-03 | ns | 1,3E-03 | 9,8E-04 | 2,9E-04 | ns |
|  |  | Bas1205T05 | 2,0E-03 | 4,3E-03 | 2,4E-03 | * | 5,9E-04 | 1,2E-03 | 2,3E-04 | ns |
|  |  | Dec8530T01 | 1,4E-02 | 0,0E+00 | 0,0E+00 | ns | 2,1E-03 | 1,5E-03 | 0,0E+00 | * |
|  |  | Bas1205T17 | 6,6E-04 | 7,6E-05 | 0,0E+00 | ns | 3,8E-04 | 1,9E-04 | 1,1E-04 | ns |
|  | Paenibacillaceae | ArcM30T12 | 0 | 0 | 0 | - | 0 | 0 | 0 | - |
|  |  | ArcP30T02 | 4,2E-02 | 5,2E-02 | 1,9E-02 | ns | 2,0E-01 | 4,3E-01 | 2,2E-01 | ns |
|  |  | ArcP30T13 | 1,5E-02 | 3,0E-02 | 6,6E-02 | * | 7,5E-02 | 1,1E-01 | 1,5E-01 | ns |
|  |  | ArcP30T20 | 5,5E-02 | 5,6E-02 | 2,4E-02 | ns | 2,3E-02 | 4,8E-02 | 1,9E-02 | ns |
|  |  | Bas0130T01 | 2,8E-04 | 1,7E-04 | 0,0E+00 | ns | 5,2E-04 | 2,2E-04 | 1,6E-04 | ns |
|  |  | Bas0130T06 | 2,4E-04 | 7,0E-04 | 0,0E+00 | * | 3,4E-03 | 2,1E-03 | 1,6E-03 | ns |
|  |  | ArcA30R01 | 1,1E-01 | 6,6E-02 | 3,2E-01 | * | 5,3E-02 | 6,8E-02 | 1,7E-01 | ns |
|  |  | ArcA30R03 | 0 | 0 | 0 | - | 0 | 0 | 0 | - |
|  |  | ArcP30T13 | 1,5E-02 | 3,0E-02 | 6,6E-02 | * | 7,5E-02 | 1,1E-01 | 1,5E-01 | ns |
|  |  | ArcA30T09 | 0 | 0 | 0 | - | 0 | 0 | 0 | - |
| Actinobacteria | Micrococcaceae | Dec8530T04 | 9,1E-01 | 3,9E-01 | 1,7E-03 | * | 8,8E-01 | 7,5E-01 | 3,2E-01 | ns |
| Proteobacteria | Moracellaceae | Dec9230T05 | 5,6E-04 | 0,0E+00 | 0,0E+00 | ns | 3,7E-01 | 3,9E-02 | 7,4E-04 | * |
|  |  | Dec9230T03 | 3,2E-04 | 0,0E+00 | 0,0E+00 | ns | 7,5E-01 | 2,6E-01 | 4,6E-03 | * |
|  |  | Dec9230T07 | 1,4E-04 | 0,0E+00 | 0,0E+00 | ns | 1,4E-01 | 1,7E-01 | 0,0E+00 | * |
|  |  | Dec9230T01 | 2,0E-04 | 0,0E+00 | 0,0E+00 | ns | 4,3E-01 | 8,8E-01 | 0,0E+00 | * |

<sup>a</sup> N / N<sub>0</sub>: N = Colony-forming units (CFU) after desiccation treatment; N<sub>0</sub> = Control group, CFU without
treatment, Signif. = comparisons were done with Kruskal-Wallis test (p<0.05, Bonferroni correction).

## Discussion

### Chloroform as a selective agent

Although, desiccation resistance accompanied the chloroform-selected strains (with some exceptions); the strains are being primarily selected by their resistance to organic solvents. The selection of desiccation resistant bacteria using organic solvents is a parallelism. The presented methodology is especially useful for its relativity quickness, and effectiveness. A comparison with a standard isolation method was made in the original report of the methodology, but studies using more traditional isolation procedures or other selective agents are encouraged. Regarding the exceptions, six of the isolated microorganisms that survived the chloroform treatment did not show any tolerance to desiccation. We hypothesize that some cells, when attached to soil particles conglomerates, may be protected against chloroform exposure and water loss. Additionally, chloroform exposure time also plays a part in the endurance of non-desiccation-tolerant microorganisms (Narvaez-Reinaldo et al., 2010). Chloroform may take more than 30 minutes to damage some cells even when they are not fully resistant to organic solvents or desiccation. Still, some isolates related to the ones presented in our work were previously described as desiccation resistant in independent experiments. In our study, we isolated the strain Dec8530T04 part of the phylum Actinobacteria and the *Micrococcaceae* family. One closely related species, *Micrococcos luteus*, was shown to be highly desiccation-tolerant (Merrick and Bruce, 1965). Another component of this phylum, *Arthrobacter siccitolerans*, was isolated by the same methodology using organic solvents as an anhydrobiosis-resistant selection method and proved highly resistant to desiccation (SantaCruz-Calvo et al., 2013). However, in a recent study, *Psychrobacter cryohalolentis K5* – isolated from a Siberian cryopeg in Russia – appeared to be especially sensitive to the desiccation process. This strain was subjected to experiments simulating conditions analogous to the Mars surface, the survival of the exposed cells were up to six orders of magnitude bellow the positive control (Smith et al., 2009). *Psychrobacter*, here isolated exclusively from Deception Island, have often been isolated from various cold environments, including ice, soil, and permafrost of Antarctica and Siberia, as well as deep-sea sediment samples (Ayala-del-Rio et al., 2010).

### Phylogeny study

The identified DNA sequences unveiled a prevalence of gram-positive isolates of the phylum *Firmicutes* within the prospected strains. This finding is consistent with the established knowledge, which indicates that these bacteria, although not part of a monophyletic group, are among the most studied bacteria that have anhydrobiosis resistance (Garcia, 2011). The main distinguishing feature of these bacteria is the presence of many layers of peptidoglycan (a polymer consisting of carbohydrates and amino acids) forming a cell wall external to the cell cytoplasmic membrane, which could favor water and small solute retention and prevent membrane collapse (Madigan et al., 2010). This phylum is also capable of forming spores (or endospores) – a dormant state that can resist for a long time to chemical and environmental challenges – allowing the organism to survive in harsh environmental conditions (Cano and Borucki, 1995). In our isolation protocol, we fixed the exposure to chloroform at 30 minutes aiming to isolate both sporulating and non-sporulating bacteria; longer exposure periods have a tendency to select only spores (Narvaez-Reinaldo et al., 2010). Although we did not directly observed spore formation, this mechanism may be responsible for the resilience of strains related to *Sporosarcina* and *Paenibacillus*. The isolation of spore-forming and non-spore-forming bacteria opens the way for future studies aiming to understand different mechanisms of resistance. Non-spore-forming bacteria have other means to protect the cells from DNA damage and protein oxidation – consequences of water loss – like the synthesis of non-reducing disaccharides, accumulation of antioxidants and expression of specialized proteins like LEA proteins, recombinases, chaperones and peroxidases (Fredrickson et al., 2008; Chakrabortee et al., 2007; Tunnacliffe et al., 2001).

For some closely related isolates, but selected from different soil samples, the desiccation tolerance results varied significantly. This could be the result of minute genetic variations leading to a considerable improvement on desiccation resistance; but also raised the question that, at least in some cases, the isolates geographical origin rather than phylogeny was determinant to desiccation tolerance. Physiological trait variations could be the result of heritable information that goes beyond DNA sequence, shaped by the physicochemical characteristics of the environment, and the root to a unique desiccation conditioning when compared to strains of different origin and the “lab-adapted” controls (Casadesús and Low, 2006).

### Desiccation medium influence

Survival rates were higher when cells were dried in the presence of the low nutrient solution R2B diluted ten times. This experiment intended to observe the influence of a low nutrient medium and its salt concentration compared to pure water in the desiccation resistance of different strains. Inorganic salts present in the formula conferred protection against osmotic stress, which might have been crucial to the survivability of strains sensitive to osmotic shock. This can explain the low survivability of the reference controls *P. halocryophilus, E. antarcticum* (both adapted to high salinity environments), when suspended in water prior to the test. All used soil samples were under the influence of the marine environment, especially in Antarctic samples collected close to the shore. Suspension in pure water added osmotic stress to the cells. Some molecules can act as stabilizer and protective agents during the anhydrobiosis state (Julca et al., 2012; Garcia, 2011). Sodium pyruvate – present in the R2B formulation and a source of energy for aerobic and anaerobic metabolisms – could also have a protective ability against reactive oxygen species (ROS) that threats desiccated cells (Giandomenico et al., 1997; Franca et al., 2007). We believe that a wider range of salt concentrations and media formulations would be necessary to understand the effect of this variable in desiccation survivability. The use of pure water in experiments seeking to understand the effect of other environmental challenges in desiccated cells may not be the best option.

### Significance to astrobiology

NASA Astrobiology Strategy regards the understanding of microbial limits of life, and its physiological adaptations, as one of the key challenges to be dealt with in astrobiology research (Hays, 2015; Des Marais et al., 2008). Within this framework, we believe the isolates described herein to be useful additions for use in future investigations to further our knowledge regarding the survival limits of life beyond Earth. In experiments seeking to understand the survivability of life to extraterrestrial conditions, extreme desiccation caused by vacuum exposure is one of the main environmental stressors. The Biomex mission, for instance, underway at the International Space Station uses the EXPOSE facility to expose microorganisms to the Low Earth Orbit and simulated Mars environmental conditions (Rabbow et al., 2012). On Earth, simulation chambers expose organisms to extreme desiccation and freeze thaw cycles in a multitude of combinations of environmental stressors (Olsson-Francis and Cockell, 2010; Lage et al., 2012). Further studies could explore the unique physiology of polar microorganisms; understand if cold survival mechanisms has a direct influence in the survivability to space environment and compare their resistance with microorganisms prone to desiccation originated from different extreme environments. The ability to withstand desiccation was even described in thermophilic microorganisms, opening the possibility to compare the convergence of adaptations in a contrasting thermal environment (Beblo et al., 2009).

In conclusion, we found that the proposed isolation methodology was able to isolate cold-adapted desiccation tolerant bacteria from assorted polar soils – with some isolates presenting a considerable resistance even after 50 days of anhydrobiosis. The solution on which the cells where dried also influenced the results; R2B medium being more protective than water. This is the first time that this methodology was applied to isolate cold-adapted, desiccation tolerant microorganisms from the Arctic and Antarctica. This study brought forward isolates of interest to future astrobiology projects seeking to understand the limits of life and looking to compare its physiology with anhydrobionts from different extreme environments.

## Acknowledgments

The present study was supported by the Brazilian agencies: National Council for Scientific and Technological Development (CNPq), Coordination for the Improvement of Higher Education Personnel (CAPES), São Paulo Research Foundation (FAPESP), Brazilian Antarctic Program (PROANTAR); and the Canadian agencies: Natural Sciences and Engineering Research Council of Canada (NSERC), Canada Research Chair program (CRC) and Canadian Foundation for Innovation (CFI).

